# Activation of T cell checkpoint pathways during β-cell antigen presentation by engineered dendritic cells promotes protection of non-obese diabetic mice from type 1 diabetes

**DOI:** 10.1101/2021.08.08.455575

**Authors:** Radhika R. Gudi, Nicolas Perez, Subha Karumuthil-Melethil, Gongbo Li, Chenthamarakshan Vasu

**Affiliations:** Department of Microbiology and Immunology, College of Medicine, Medical University of South Carolina, Charleston, SC-29425; Department of Surgery, College of Medicine, University of Illinois, Chicago, IL 60612

**Keywords:** Type 1 diabetes, autoimmunity, dendritic cells, T cell repressor/inhibitory receptors, beta-cell antigen, T cell tolerance

## Abstract

Defective immune regulation has been recognized in type 1 diabetes (T1D). Immune regulatory T cell check-point receptors, which are generally upregulated on activated T cells, have been the molecules of attention as therapeutic targets for enhancing immune response in tumor therapy. We reported that dendritic cells (DCs) that are engineered to express selective ligands for checkpoint receptors can induce effective T cell tolerance in antigen-immunization models. Here, we show that pancreatic β -cell antigen (BcAg) presentation by engineered tolerogenic-DCs (tDCs) that express CTLA4 selective ligand (B7.1wa) or a combination of CTLA4, PD1 and BTLA selective ligands (B7.1wa, PD-L1, and HVEM-CRD1 respectively; multiligand-DCs) causes an increase in regulatory cytokine and T cell (Treg) responses and suppression of the effector T cell function as compared to engineered control-DCs. Non-obese diabetic (NOD) mice treated with BcAg-pulsed CTLA4-ligand-DCs and multiligand-DCs at pre-diabetic and early-hyperglycmic stages showed significantly lower degree of insulitis, higher frequencies of insulin-positive islets, profound delay in, and reversal of, hyperglycemia for a significant duration. Immune cells from the tDC treated mice not only produced lower amounts of IFNγ and higher amounts of IL10 and TGFβ1 upon BcAg challenge, but also failed to induce hyperglycemia upon adoptive transfer. While both CTLA4-ligand-DCs and multiligand-DCs were effective in inducing tolerance, multiligand-DC treatment produced an overall higher suppressive effect on effector T cell function and disease outcome. These studies show that enhanced engagement of T cell checkpoint receptors during BcAg presentation can modulate T cell function and suppress autoimmunity and progression of the disease in T1D.

## Introduction

Type 1 diabetes (T1D) is an autoimmune disorder resulting from the progressive destruction of insulin-producing pancreatic β cells by autoreactive lymphocytes, leading to insulin deficiency and hyperglycemia. Pancreatic islet β-cell antigen (BcAg)-specific T cells play a major role in the disease process, and they arise and expand under genetic susceptibility due to defective immune regulation^1^. Immune regulatory defects and disease susceptibility in both humans and experimental animals have been linked to many genetic factors including the loci encoding for T cell inhibitor/repressor receptors such as Cytotoxic T-lymphocyte-associated antigen 4 (CTLA4)^2,3^. Hence, approaches targeting these defects as well as enhancing the T cell repressor function could be valuable for T1D therapy.

The inhibitory/repressor receptors, also referred to as checkpoint receptors, expressed on T cells are critical for preventing and regulating self-antigen specific immune responses^4-6^. CTLA4, programmed cell death-1 (PD1) and B- and T-lymphocyte attenuator (BTLA) are the major checkpoint receptors of CD28 family expressed on T cells^7,8^. Surface expression levels of these receptors are upregulated upon activation of T cells as a homeostatic mechanism. While CTLA4 down-regulates the T cell response upon binding to CD80 and CD86 of antigen presenting cells (APCs), these interactions also curb activation of APCs through reverse signaling^9,10^. CTLA4 is a master regulator of peripheral T cell tolerance, T cell homeostasis and regulatory T cell (Treg) function^11-21^. CTLA4 deficiency causes massive lymphoproliferative condition in mice, leading to their death at as early as 4 weeks of age^21^. PD1 is induced on T and B cells upon activation and its interaction with PD-L1 and PD-L2 activates negative regulatory function^22,23^. PD1 deficiency leads to increased susceptibility to autoimmunity^24^. BTLA is expressed on activated T and B cells, and its signals can also suppress the T-cell response^8^. BTLA interacts with the TNFR family member, HVEM (a herpes virus entry mediator)^25,26^. While HVEM also interacts with T cell activation receptor LIGHT, its interaction with BTLA and CD160 on activated T cells through its CRD1 region induces an inhibitory response in T cells^26,27^. Considering their role in immune tolerance, T cell checkpoint pathways, initiated by CTLA4 and PD1 particularly, are being targeted to treat cancers^28-30^. However, these therapies also produced autoimmune etiology in patients^31-34^, further substantiating their role in maintaining peripheral self-antigen tolerance.

Previously, we have shown that the dominant engagement of CTLA-4 on T cells from target tissues and antigen presenting dendritic cells (DCs) can induce Treg responses, antigen specific T cell tolerance, and prevent and suppress experimental autoimmune thyroiditis (EAT) and spontaneous type 1 diabetes (T1D) in mouse models^35-39^. Recently, we tested if exogenous expression of checkpoint-receptor selective ligands in DCs for enhancing T cell inhibitory signals can generate tolerogenic antigen presenting cells (tDCs/tAPCs)^40^. DCs that exogenously express selective ligands for CTLA4, PD1 and BTLA individually (monoligand-DCs) or in combination (multiligand-DCs) induced an antigen specific immune tolerance. These engineered DCs, CTLA4-ligand-DCs and multiligand-DCs particularly, can induce profound inhibition of T cell proliferation, modulation of cytokine response, generation of T cells with a regulatory phenotype and suppress autoimmunity in a thyroglobulin immunization induced EAT model. However, whether this engineered DC approach can be employed to suppress spontaneous autoimmunity is not known. Here, we show that engineered CTLA4-ligand-DCs and multiligand-DCs that are pulsed with pancreatic β-cell antigen peptide cocktail (BcAg) induced immune regulatory cytokine and Treg responses and prevented T1D in non-obese diabetic (NOD) mice for a profound duration, when treated at pre-diabetic stage. Furthermore, early-hyperglycemic mice that received these tDCs showed reversal of hyperglycemia for a significant duration. Overall, this study demonstrates that enhancing checkpoint receptor signaling during self-antigen presentation by DCs can effectively suppress autoimmunity and progression of the disease in T1D.

## Materials and methods

### Mice, antibodies, antigens, and other reagents

NOD/Ltj and NOD-BDC2.5 TCR-transgenic (BDC2.5 mice) were originally purchased from the Jackson laboratory (Bar Harbor, ME, USA). Foxp3-GFP-knockin (ki)^41^ mice in the B6 background were kindly provided by Dr. Vijay Kuchroo (Harvard Medical School, MA). NOD-Foxp3-GFP-ki mice and NOD-BDC2.5-Foxp3-GFP-ki mice were generated as described in our previous reports^42,43^. Breeding colonies of these mice were established and maintained in the specific pathogen free facility of the Medical University of South Carolina (MUSC). Glucose levels in the tail vein blood samples of mice were monitored with the Ascensia Microfill blood glucose test strips and an Ascensia Contour blood glucose meter (Bayer, Pittsburgh, PA). The mice with glucose levels >250 mg/dl for two consecutive bleeds were considered diabetic. Mice with blood glucose levels: 140-250 mg/dl and >450 mg/dl were considered early hyperglycemic and overt-diabetic respectively. All experimental protocols were approved by the Institutional Animal Care and Use Committee (IACUC) of MUSC. All methods in live animals were carried out in accordance with relevant guidelines and regulations of this committee.

Immunodominant β-cell-Ag (BcAg) peptides [namely: 1) insulin B(9-23); 2) GAD65(206-220); 3) GAD65(524-543); 4) IA-2β(755-777); 5) IGRP(123-145), 6) GAD65(286-300), 7) Insulin B(15-23), and 8) IGRP(206-214)], and BDC2.5 TCR-reactive peptide (YVRPLWVRME; referred to as BDC2.5-peptide) were used. Peptides 1-8 were pooled at an equal molar ratio and used as BcAg to pulse the DCs for in vitro and in vivo experiments, as done in our earlier studies, using some of these peptides^37,39,43,44^.

### Generation of cDNA vectors and lentivirus production

The third-generation replication-incompetent lentiviral cDNA cloning vector and packaging plasmids were purchased from System Bio Inc and modified^40^. Details on the generation of vector constructs: pCDH1 vector with cDNA for B7.1wa, PD-L1 or HVEM-CRD1 and GFP reporter under a separate promoter are described in our previous report^40^. Most experiments employed DCs engineered using these constructs. pCDH1-vector encoding a “multi-gene/cDNA sequence” (B7.1wa, HVEM-CRD1 and PDL1 cDNAs separated by self-cleaving P2A and T2A peptide sequences: pCDH1-EF1-copGFP-B7.1wa-T2A-HVEM-CRD1-P2A-PDL1; pCDH1-multiligand vector) (shown in supplemental Fig. 1) was also constructed by gene synthesis and custom cloning service of Genscript and used in some experiments where specified. Comparable results were obtained when single-cDNA and multi-cDNA constructs were used in in vitro assays. For lentivirus production, HEK293T cells were transfected with cDNA expression vectors along with packaging vectors by calcium phosphate method. GPRG cells^45^, provided by National Gene Vector Biorepository (Indianapolis, IN, USA), were also used as virus packaging cells in some experiments. Preparation and titration of virus particles are described in our recent report^40^.

### Generation of engineered BMDCs

Generation of DCs from BM cells (BMDCs) and engineering using lentiviral particles are described in our previous reports^37-40^. Briefly, BM cells from prediabetic NOD mice were cultured in complete RPMI 1640 medium containing 10% FBS, in the presence of GM-CSF (20 ng/ml), at 37°C in 5% CO_2_ for 3 days and then for further 3 days in fresh complete RPMI 1640 medium, containing GM-CSF (20 ng/ml) and IL-4 (5 ng/ml). For lentiviral transduction, cells from 3-day-old cultures were used. A final concentration of >5×10^8^ TU/ml virus (for individual ligand-DCs and control-DCs) or a cocktail of >1.5 ×10^9^ TU/ml virus (for multiligand-DCs) was used for transducing cells in the presence of polybrene (8 μg/ml) and protamine sulfate (10 μg/ml) for up to 24h, washed and cultured in fresh medium containing GM-CSF and IL4 for an additional 48h. Transduction efficiency was examined by microscopy and/or FACS, and the cells that showed transduction efficiency of >90% were washed thoroughly and counted before in vitro or in vivo use. Ligand functionality was determined by assessing the increase in soluble receptor binding by ligand-DCs as compared to control-DCs having endogenous ligand expression.

### In vitro antigen presentation assay

BDC2.5-peptide pulsed NOD BMDCs (2 × 10^4^ cells/well) were plated in triplicate in 96-well U-bottom tissue culture plates along with unlabeled or CFSE-labeled purified CD4+ T cells from NOD-BDC2.5 or NOD-BDC2.5-Foxp3-GFP mice (1 × 10^5^ cells/well) in complete RPMI 1640 medium. After 4 days of culture, cells were subjected to surface and/intracellular staining using fluorochrome-labeled Abs and examined for CFSE dilution or cytokine production by FACS. Cells were subjected to brief activation using PMA and ionomycin (4h) in the presence of Brefeldin A, before staining to detect intracellular cytokines. In some assays, equal number of live cells from primary cultures were seeded along with BDC2.5 peptide and freshly isolated NOD-mouse splenic DCs for 24 h, and the spent media were tested for cytokine levels by ELISA or Luminex multiplex assay. In some assays, BcAg (peptide cocktail) pulsed DCs were incubated with T cells from the pancreatic lymph nodes (PnLNs) of early-hyperglycemic mice for up to 4 days and subjected to various assays.

### Treatment of NOD mice with engineered DCs

Female NOD mice of different age groups and disease stages were injected i.v. with 2×10^6^ BcAg-pulsed control-DCs or ligand-DCs (B7.1wa-DCs or multiligand-DCs) once or twice, at a 15-day interval. Cohorts of animals were examined for blood glucose levels every week. In some experiments, two or four-weeks post-treatment, mice were euthanized to test for T cell response to ex vivo challenge with the BcAg, T cell phenotype, and/or insulitis.

### Adoptive transfer of immune cells to NOD-Rag1-/- mice

Total PnLN cells of engineered DC treated mice were cultured overnight in round-bottom 96-well plates in the presence of anti-CD3-Ab (2 μg/ml), washed, and injected into 6-8-week-old NOD-*Rag1-/-* mice. In some experiments, splenic T cells enriched from the spleens of untreated early-hyperglycemic mice were cultured in the presence of BcAg pulsed DCs for 4 days, and live T cells enriched from these primary cultures were injected (1 × 10^6^ cells/mouse) i.v. into NOD-*Rag1-/-* mice. These T cell recipients were tested for blood glucose levels every week. In some experiments, purified CD4+ T cells from 4-week-old BDC2.5 TCR-Tg mice that were cultured in the presence of BDC2.5 peptide-pulsed various engineered DCs, as described above and cells from these cultures (1 × 10^6^ cells/mouse) were injected i.v. into 4-week-old male WT-NOD mice. Blood glucose levels in these mice were tested every other day. Cohorts of mice were euthanized at different time-points to determine the degree of insulitis.

### Histochemical analysis of pancreatic tissues

Pancreatic tissues were fixed in 10% formaldehyde and 5-μm paraffin sections were made and stained with H&E. Stained sections were analyzed in a blinded fashion using a grading system: 0, no evidence of infiltration; 1, peri-islet infiltration (<5% of the islet area); 2, <25% infiltration; 3, 25-50% infiltration; and 4, >50% infiltration as described in our earlier studies^37,39,43,44,46,47^. In some experiments, pancreatic sections were stained using anti-insulin antibody followed by Alexa fluor 488- or 568-linked secondary antibodies and DAPI and scored for insulitis based on DAPI-positive cells in islet areas and insulin expression. Insulitis was scored as described for H&E stained sections and insulin positive and negative islets were counted as described in our previous reports^42-44^.

### Statistical analysis

Mean, SD, and statistical significance (*p*-value) were calculated using Microsoft Excel or GraphPad prism statistical application. In most cases, values of the individual test group (ligand-DC group) were compared with that of control group (control-DC group). For most comparisons, Mann-Whitney or paired *t*-test was employed. Statistical analysis of insulitis severity was done by Fisher’s exact test. T1D incidence in different groups was compared by employing log-rank test. Specific methods used for obtaining *p*-values were described under figure legends. *p* ≤ 0.05 was considered significant.

## Results

### T cell-checkpoint receptor selective ligand expressing DCs modulate T cell response in vitro

In a recent report, we showed that exogenous expression/overexpression of selective ligands of CTLA4, PD1 and BTLA receptors could make the DCs produce, upon antigen presentation, tolerogenic effects on T cells, presumably through enhanced receptor engagement^40^. We engineered the BMDCs to express B7.1wa, PD-L1 and HVEM-CRD1 exogenously, individually and in combination and characterized their receptor-binding, antigen presenting and tolerogenic properties extensively^40^. WT B7.1 binds to both CTLA4 and CD28, and WT HVEM binds to BTLA and LIGHT. CTLA4- and BTLA-selective binding abilities of B7.1wa and HVEM-CRD1 were described before^48,49^. We found that B7.1wa-DCs and multiligand-DCs have a better ability to induce robust Treg responses and overall immune modulation compared to control-DCs, and other mono-ligand DCs such as PD-L1-DCs and HVEM-CRD1-DCs^40^. This report which used a mouse thyroglobulin immunization model showed that B7.1wa-DCs and multiligand-DCs are effective in modulating T cell function and suppressing autoimmune thyroiditis. However, whether such engineered DCs are useful for modulating spontaneous autoimmunity such as T1D is not known. Considering the superior tolerogenic effects of B7.1wa-DCs and multiligand-DCs, the current study examined the antigen presenting and tolerogenic properties of these engineered-DCs **(Fig. 1A & B)** in the context of T1D in NOD mice.

**FIGURE 1:**
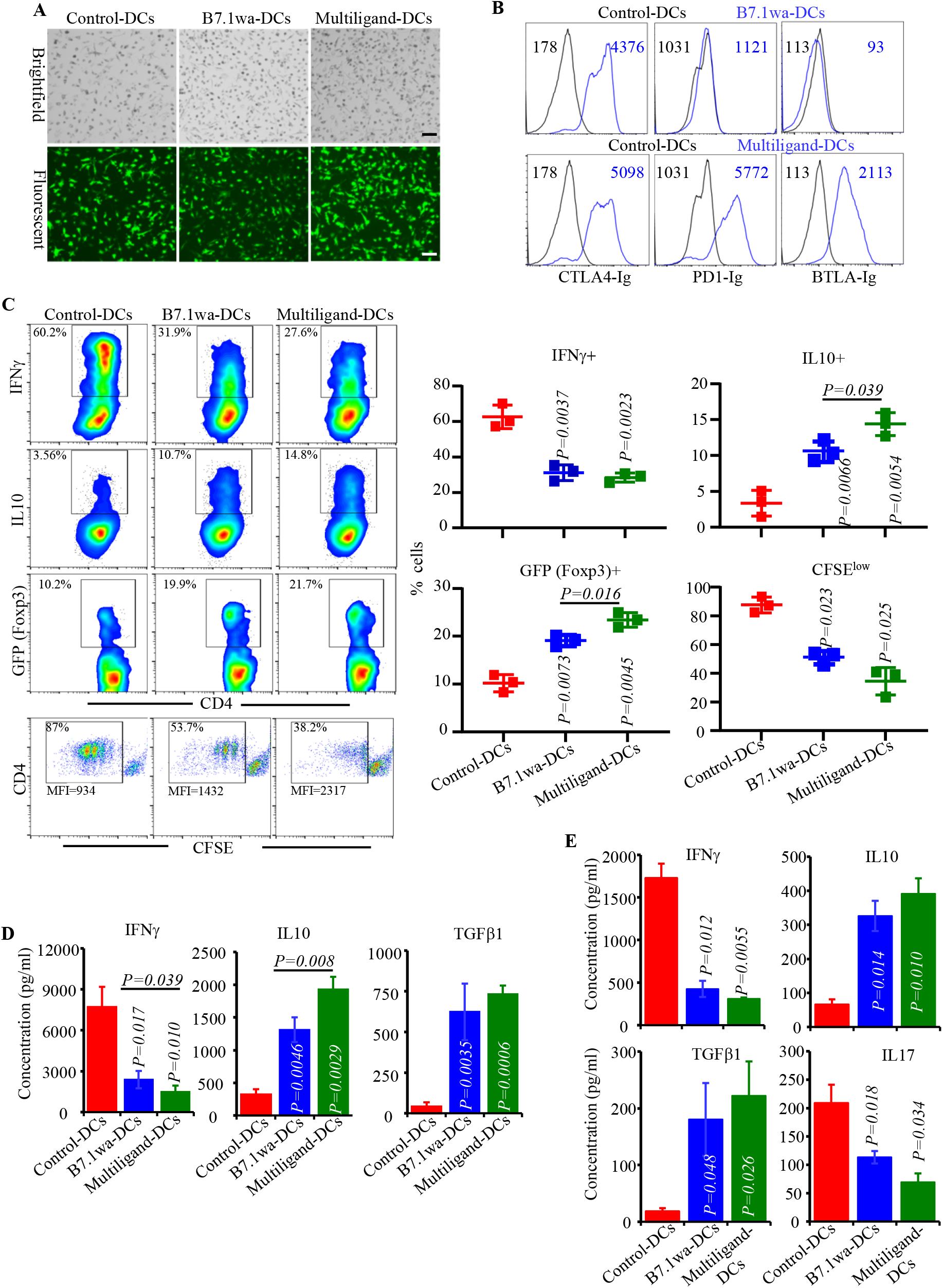
Characterization of the antigen presenting properties of engineered NOD mouse DCs. NOD mouse BM DCs were transduced with lentiviral vectors as described in Materials and methods. **A)** Examples of transduction of DCs using various preparations of virus with GFP reporter are shown. **B)** Functional ligand expression levels on DCs were determined after incubating with soluble receptors (CTLA4-Ig, PD1-Ig or BTLA-Ig), followed by PE-labeled Fab(2) fragment of anti-IgG (Fc specific) Ab and testing by FACS. Histogram overlay graphs (representative of 3 independent experiments) showing ligand specific staining of control virus transduced and ligand virus transduced DCs along with mean fluorescence intensity (MFI) values are shown. Both control and ligand DCs were also stained using control Ig reagents to assess the background staining (not shown). **C)** BDC2.5-peptide pulsed engineered DCs were used in antigen presentation assays by culturing them with CD4+ cells isolated from BDC2.5 mouse spleens. After 4 days, cells from the primary cultures were stimulated using PMA and ionomycin, in the presence of Brefeldin A for 4h, stained for intracellular cytokines IFNγ and IL10, and subjected to FACS analysis. CD4+ T cells from BDC2.5-Foxp3-GFP mice were also used in primary cultures for 4 days and subjected to FACS to determine GFP+ CD4+ cell frequencies by FACS. In addition, CFSE labeled CD4+ T cells from BDC2.5 mice were used in primary culture for 4 days and examined for percentage of cells with CFSE dilution and their MFI by FACS. CD4+ cells were gated and representative FACS plots (left) and Mean±SD values of 3 independent experiments (transduction using three preparations of lentivirus), each done in triplicate (right), are shown. **D)** Equal number of BDC2.5 T cells isolated from primary cultures were cultured with BDC2.5 peptide pulsed fresh splenic DCs for 48h, and supernatants were subjected to Luminex multiplex assay or ELISA. **E)** PnLN cells from early hyperglycemic NOD mice were cultured with BcAg-pulsed engineered-DCs for 4 days, equal number of T cells isolated from these primary cultures were challenged using BcAg-pulsed fresh splenic DCs, and supernatants were tested for cytokine levels as done for BDC2.5 T cells. pCDH1-multiligand vector, instead of individual ligand vectors, was used for generating multiligand-DCs for panel E. Mean± SD values of 3 independent experiments (transduction using three preparations of lentivirus), each done in triplicate, are shown for panels D and E. *P*-value by paired *t*-test for panels C-E. All *P*-values are in comparison with control-DC group unless indicated (for B7.1wa group vs multiligand-DC group).

To assess the abilities of inhibitory ligand expressing NOD-mouse DCs to activate T cells, control-DCs, B7.1wa-DCs and multiligand-DCs were pulsed with BDC2.5 peptide and cultured in the presence of CD4+ T cells enriched from NOD-BDC2.5 TCR-Tg mice or NOD-BDC2.5-Foxp3-GFP-ki mice. As observed in **Fig. 1C**, T cells cultured with B7.1wa-DCs and multiligand-DCs showed lower frequencies of IFNγ+ cells, higher frequencies of IL10+ and Foxp3+ cells, and a lower rate of proliferation as compared to those cultured with control-DCs. Ligand-DC activated BDC2.5 T cells produced higher amounts of immune regulatory cytokines IL10 and TGFβ1 and showed diminished production of the pro-inflammatory cytokine IFNγ upon rechallenge with BDC2.5 peptide **(Fig. 1D)**. We then examined the ability of BcAg pulsed engineered DCs to modulate the properties of splenic T cells obtained from early-hyperglycemic WT NOD mice in vitro. T cells from these primary cultures were challenged with BcAg and examined for the levels of secreted cytokines. As observed in **Fig. 1E**, Ag presentation by B7.1wa and multi-ligand DCs resulted in the modulation of T cell phenotype, as indicated by significantly lower levels of proinflammatory IFN-γ and IL17 and higher immune regulatory IL-10 and TGF-β1 cytokine responses, compared to T cells activated using control-DCs. These results show that enhanced DC-directed negative signaling, during BcAg presentation, suppresses the pro-inflammatory activation of, and promotes immune regulatory features in, T cells.

### B7.1wa-DC and multiligand-DC activated T cells fail to induce hyperglycemia

To assess if the in vitro activation of T cells by ligand-DCs has an impact on their diabetogenic properties, WT-NOD and BDC2.5 T cells were cultured in the presence of control-DCs, B7.1wa-DCs and multiligand-DCs in vitro for 4 days as described for Fig. 1, equal number of enriched live cells from these cultures were injected i.v. into NOD-*Rag1-/-* mice or young WT NOD mice and monitored for hyperglycemia. **Fig. 2A** shows that 100% of NOD-*Rag1-/-* mice that received control-DC activated WT T cells developed hyperglycemia during 8-12-week period after cell transfer. However, mice that received B7.1wa-DC and multiligand-DC activated WT T cells failed to develop hyperglycemia, at least for 18 weeks post-cell transfer. Pancreatic tissues of B7.1wa-DC and multiligand-DC activated T cell recipients that were euthanized 4-weeks post cell-transfer, showed significantly lower degree of insulitis compared to those which received control-DC activated T cells **(Fig. 2B)**. In BDC2.5 T cell recipient WT NOD mice, while 100% of mice that received control-DC activated T cells developed overt-hyperglycemia within 8 days, recipients of B7.1wa-DC and multiligand-DC activated T cells failed to show hyperglycemia for at least 15 days **(Fig. 2C)**. Further, recipients of ligand-DC activated T cells showed little or no insulitis on day 5 post-cell transfer as opposed to 100% of the islets of control group showing severe grade 3 or grade 4 insulitis **(Fig. 2D)**. Overall, these results show that T cell inhibitory ligand expressing-DCs can suppress the diabetogenic/pathogenic activation of T cells upon antigen presentation in vitro.

**FIGURE 2:**
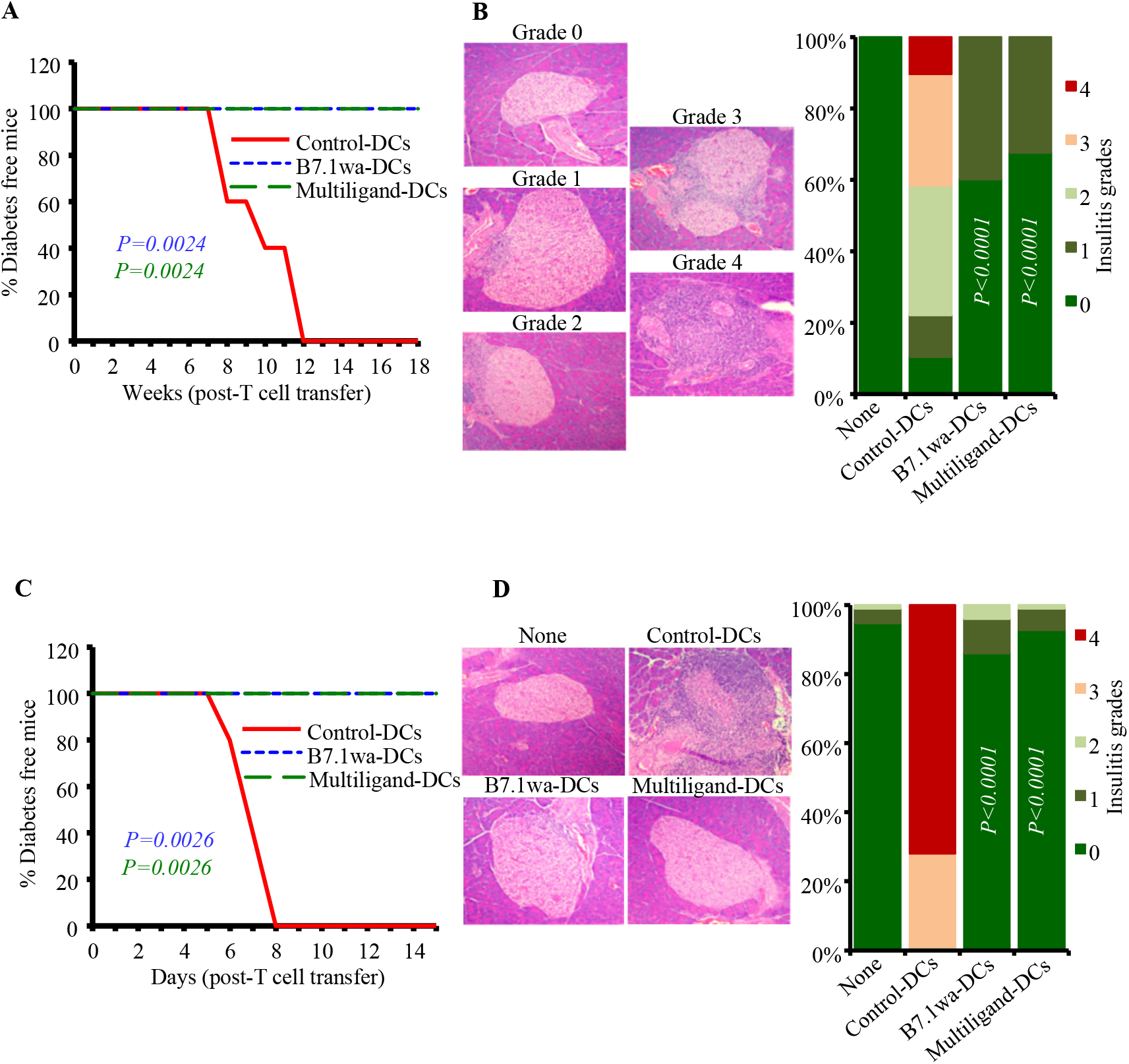
Diabetogenic properties of T cells activated in vitro using engineered-DCs. PnLN T cells from early-hyperglycemic WT-NOD mice and CD4+ T cells from BDC2.5 mice were subjected to primary culture in the presence of various DC preparations for 4 days as described for Fig. 1. **A&B)** Live T cells from primary cultures of WT PnLN T cells were enriched and injected into 6-week-old NOD-*Rag1*-/- mice (1×10^6^ cells/mouse) and tested for blood glucose levels every week for up to 18 weeks (A; 5 mice/group) or euthanized 30 days post-cell transfer (B; 4 mice/group). pCDH1-multiligand vector, instead of individual ligand vectors, was used for generating multiligand-DCs for panels A&B. **C&D)** Live T cells from the primary cultures of BDC2.5 cells were injected into 4-week-old WT male NOD mice and tested for blood glucose levels every week for up to 15 weeks (C; 5 mice/group) or euthanized 5-days post-cell transfer (D; 4 mice/group). At least 120 islets from intermittent sections were examined for each group (at least 30 islet areas/mouse) for panels B and D. *P*-values by log-rank test for panels A and C, and Fisher’s exact test for panels B and D. All *P*-values are in comparison with control-DC group. Group “none” did not receive T cells.

### Single injection of B7.1wa-DCs or multiligand DCs causes significant modulation of BcAg specific T cell response in NOD mice

To determine the effect of treatment with engineered DCs on immune cells, ten-week-old pre-diabetic female NOD mice were injected once with control-DCs, B7.1wa-DCs and multiligand-DCs, euthanized after 15 days, and PnLN cells were examined for their proliferative response upon challenge with BcAg, and cytokine and Foxp3 expressing CD4+ T cell frequencies. As observed in **Fig. 3A**, PnLN T cells from B7.1wa-DC and multiligand-DC recipients proliferated at significantly lower rate compared to the cells from control-DC recipient mice. Intracellular staining of fresh PnLN cells showed that while IFNγ+ and IL17+ T cell frequencies were significantly lower in B7.1wa-DC and multiligand-DC recipients, IL10+ and Foxp3+ T cell frequencies were higher in these mice compared to control-DC recipients.

**FIGURE 3:**
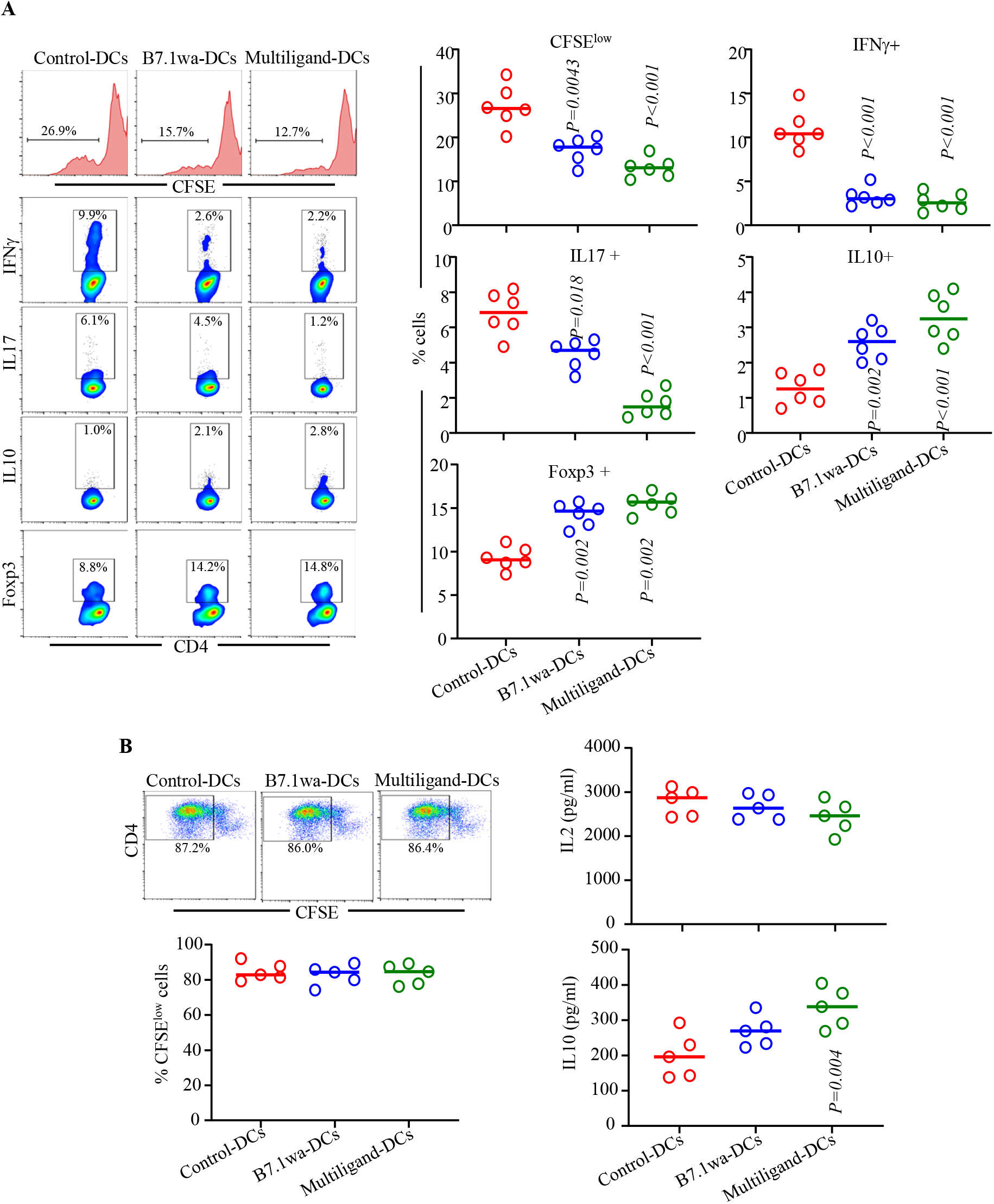
Treatments using B7.1wa-DCs and multiligand-DCs result in modulation of immune response to BcAg in vivo. BcAg-pulsed engineered DCs were injected once into 10-week-old pre-diabetic NOD mice (2×10^6^ cells/mouse; single injection). **A)** Cohorts of mice (6 mice/group) were euthanized 15-days post-treatment, PnLN cells were labeled with CFSE, cultured in the presence of BcAg for 72h and examined for CFSE dilution by FACS. Fresh PnLN cells were also cultured in the presence of PMA/Ionomycin and Brefeldin A for 4h and subjected to intracellular staining and FACS to detect cytokine positive cell frequencies. In addition, fresh cells were examined for Foxp3+ cell frequencies by FACS. CD4+ T cells were gated for these analyses. **B)** Spleen cells from additional cohorts of mice (5 mice/group) were labeled using CFSE, stimulated using anti-CD3 antibody (2 μg/ml) for 72h, and CFSE dilution was examined by FACS. Supernatants from these cultures were tested for secreted cytokines by multiplex assay. *P*-value by Mann-Whitney test. All *P*-values are in comparison with control-DC group.

To assess if BcAg loaded ligand-DC treatment, compared to control-DC treatment, impacts the global T cell function, spleen cells from the treated mice were examined for proliferative, IL2 and IL10 responses upon stimulation using anti-CD3 antibody. **Fig. 3B** shows that the degree of anti-CD3 antibody induced proliferation of CD4+ T cells was not significantly different between control-DC and ligand-DC treated mice. However, modestly lower IL2 responses, albeit not significant statistically, were observed with splenic T cells from ligand-DC treated mice compared to control-DC recipients. On the other hand, immune regulatory cytokine, IL10 was produced at higher levels by cells from ligand-DC treated mice, multiligand-DC recipients particularly, compared to control-DC recipients. Overall, sustained ability of most T cells to proliferate and produce comparable amounts of IL2 upon CD3 stimulation suggests that BcAg-pulsed ligand-DC induced modulation of immune response is primarily BcAg specific and immune regulatory in nature.

### B7.1wa-DC and multiligand-DC treatments, at pre-diabetic stage, exert profound protection from T1D and immune modulation

To assess the impact of treatment using engineered DCs on T1D incidence in NOD mice, ten-week-old pre-diabetic female NOD mice were injected i.v., twice at 15-day interval, with BcAg-pulsed control-DCs, B7.1wa-DCs or multiligand-DCs and monitored for hyperglycemia. As shown in **Fig 4A**, diabetes onset was detected in the control groups of mice within 4-6 weeks of monitoring as compared to 18-weeks post-treatment in multi-ligand DC treated mice and 12 weeks post-treatment in B7.1wa-DC treated mice. While about 60% and more than 80% respectively of B7.1wa-DC and multiligand-DC treated mice remained euglycemic for at least 30-weeks post treatment initiation, 100% of the control-DC recipient mice turned hyperglycemic during this period. To assess if treatment using engineered-DCs causes protection of pancreatic islets from destruction, tissues from treated mice were examined for insulitis, 30 days post treatment. **Fig. 4B** shows that mice that received B7.1wa-DCs and multiligand-DCs had significantly lower frequencies of islets with severe insulitis compared to control-DC recipients. While about 70% of pancreatic islets from control-DC recipients showed severe insulitis grade of ≥3, only about 30% and 10% of pancreatic islets in mice that received B7.1wa-DCs or multiligand-DCs respectively, had severe insulitis. Overall, these results show that treatment using BcAg-presenting, T cell checkpoint-receptor ligand DCs, multiligand-DCs particularly, can prevent hyperglycemia for a profound duration.

**Figure 4:**
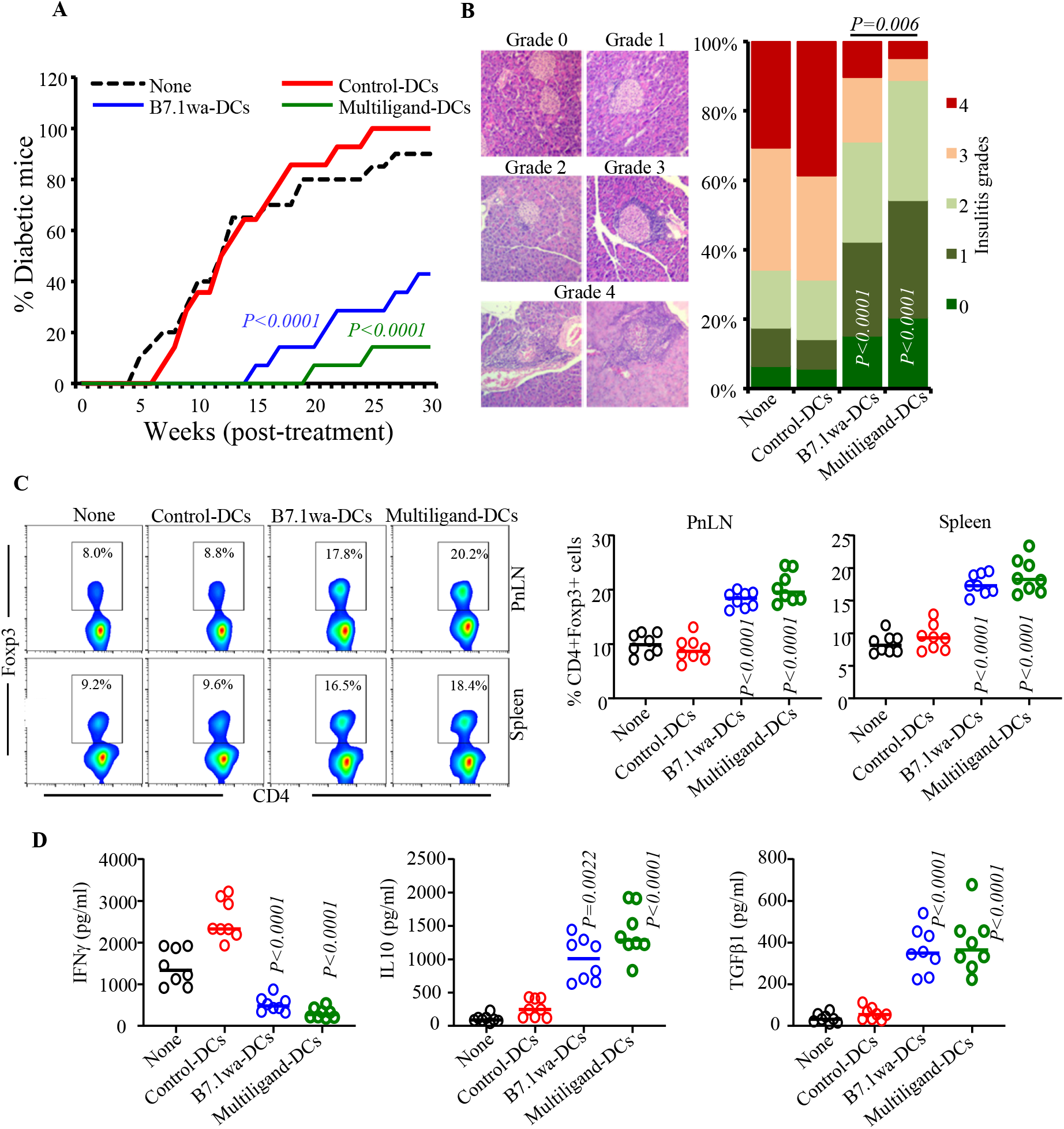
B7.1wa-DC and multiligand-DC treated pre-diabetic NOD mice show significant delay in disease onset. Ten-week-old pre-diabetic NOD mice were left untreated (none) or injected i.v. with engineered-DCs twice at a 15-day interval (2×10^6^ cells/mouse/injection). **A)** Cohorts of mice (14 mice/group for treated groups and 20 mice for “none” group) were monitored for hyperglycemia by testing for blood glucose levels every week. **B)** Cohorts of mice (5 mice/group) were euthanized 30 days post-treatment, pancreatic sections were subjected to H&E staining and grading for insulitis severity. At least 150 islet areas from intermittent sections were examined for each group (at least 30 islet areas/mouse). **C)** Cohorts of mice (8 mice/group) were euthanized 15-days post-treatment, PnLN and spleen cells were stained for Foxp3 and analyzed by FACS. **D)** PnLN cells were also cultured in the presence of BcAg for 72h and supernatants were tested for secreted cytokines by multiplex assay or ELISA. *P*-value by log-rank test for panel A, Fisher’s exact test for panel B, and Mann-Whitney test for panels C and D. All *P*-values are in comparison with control-DC group unless indicated for B7.1wa group vs multiligand-DC group.

Fifteen days post-treatment, cohorts of mice were examined for Foxp3+ T cell frequencies and BcAg-challenge induced cytokine secretion. As observed in **Fig. 4C**, Foxp3+ T cell frequencies were profoundly higher in the spleen and PnLN of B7.1wa-DC and multiligand-DC treated mice compared to control-DC treated mice. PnLN cells from B7.1wa-DC and multiligand-DC treated mice, compared to control-DC recipients, showed significantly lower IFNγ, and higher IL10 and TGFβ1 secretion upon ex vivo challenge exposure to BcAg **(Fig. 4D)**. These results suggest that the B7.1wa-DC and multiligand-DC treatments promote protection of β-cells from destruction, perhaps through enhancing immune regulation, leading to prevention of hyperglycemia.

### Treatment with B7.1wa-DCs or multiligand-DCs at early-hyperglycemic stage results in transient reversal of hyperglycemia

To determine if blood glucose levels in mice with early-hyperglycemia can be altered by treating with ligand-DCs, female-NOD mice with blood glucose levels between 140-250 mg/dl (12-25 weeks of age) were injected twice, 15-days apart, with control-DCs, B7.1wa-DCs or multiligand-DCs and examined for blood glucose levels every week. As shown in **Fig. 5A**, while 100% of the control-DC treated mice progressed to overt hyperglycemia within 8 weeks, all mice that received ligand-DCs reverted to and remained euglycemic for at least 8 weeks post-treatment initiation. Thirty days post-treatment (45 days post-treatment initiation), pancreatic tissues were collected from cohorts of mice to examine for insulitis and to assess the abundance of insulin positive islets. Tissue sections were stained using anti-insulin and anti-glucagon antibodies and examined for the frequencies of “functional islets”. As shown in **Fig. 5B**, about 80% and 90% of islet areas, with or without immune cell infiltration, of B7.1wa-DC and multiligand-DC recipient mice respectively showed some degree of islet function, as indicated by both insulin and glucagon positivity, compared to about 5% insulin positive islet remnants in control-DC treated mice. These results show that reversal of hyperglycemia is possible by treatment with T cell-checkpoint receptor ligand expressing engineered DCs.

**Figure 5:**
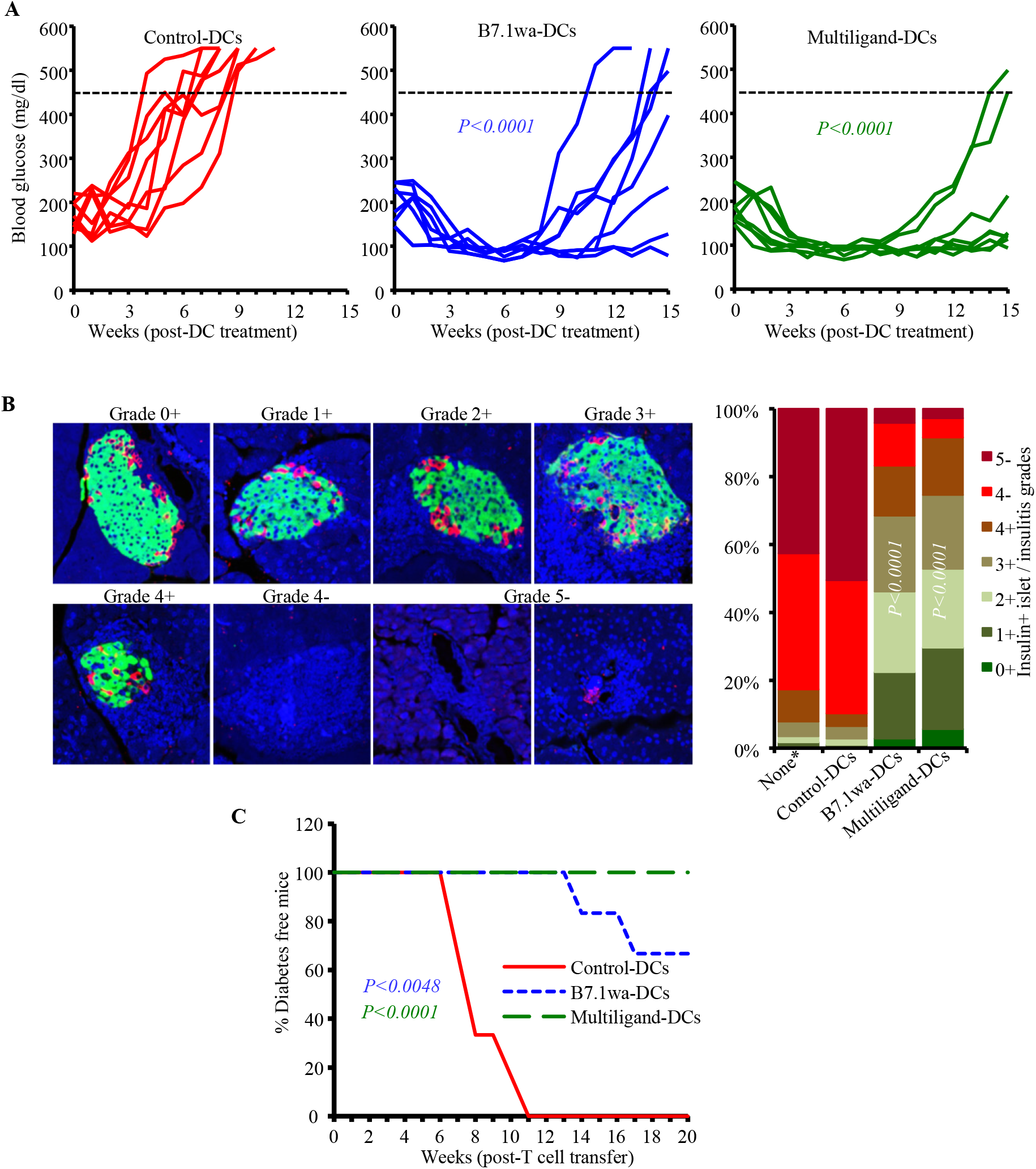
B7.1wa-DC and multiligand-DC treatments at early-hyperglycemic stage results in reversal of hyperglycemia for significant durations. Early-hyperglycemic female NOD mice (blood glucose: 140-250 mg/dl) were treated with control-DCs and B7.1wa-DCs and multiligand-DCs twice as described for Fig. 4. pCDH1-multiligand vector, instead of individual ligand vectors, was used for generating multiligand-DCs in this experiment. **A)** Blood glucose levels were examined every week and glucose values of individual mice are shown. Dotted line represents overt-hyperglycemic (blood glucose: >450 mg/dl). **B)** Cohorts of mice were euthanized 30-days post treatment, and pancreatic sections were subjected to insulin (green), glucagon (red) and DAPI (blue) staining and insulin positive islets and immune cell infiltration, based on DAPI staining, were scored. *Pancreatic tissues from untreated mice that were at early-hyperglycemic stage at the time of euthanasia were included as “none” control group to assess the pre-treatment frequency of insulin positive islets. **C)** PnLN cells from individual mice (6 mice/group), cultured for 24h with BcAg, and injected (i.v.) into NOD-*Rag1-/-* mice (about 1×10^6^ cells/mouse; one recipient/donor) and examined for blood glucose levels every week, for up to 20 weeks. *P*-value by log-rank test with overt hyperglycemia for two consecutive weeks as the end-point for panel A, Fisher’s exact test for comparing grades ≤3+ and ≥4+ among groups for panel B, and log-rank test for panel C. All *P*-values are in comparison with control-DC group.

As observed in prediabetic mice (Fig. 4C and 4D), PnLN cells from these early-hyperglycemic groups of mice showed significantly higher frequencies of Foxp3+ T cells and higher production of IL10 and TGFβ and lower secretion of IFNγ compared to control-DC recipients (not shown). To determine the pathogenic effect of these immune cells, PnLN cells from DC treated individual mice (harvested 30-days post-treatment) were cultured in the presence of anti-CD3 antibody for 24h, injected into NOD-*Rag1-/-* mice and monitored for hyperglycemia. As observed in **Fig. 5C**, all recipients of PnLN cells from control-DC treated donors turned diabetic within 11 weeks of cell transfer. However, mice that received PnLN cells from B7.1wa-DC and multiligand-DC treated mice showed a significant delay in the onset of hyperglycemia, or failed to develop hyperglycemia, during the 20-week monitoring period. This suggested that the diabetogenic properties of immune cells of early-hyperglycemic mice were profoundly suppressed in ligand-DCs treated mice. Overall, these results show that T cell checkpoint-receptor selective ligand expressing DCs, multiligand-DCs particularly, can be effective tolerogenic DCs and used to induce self-antigen specific T cell immune modulation for preventing and suppressing spontaneous autoimmunity in T1D

## Discussion

BcAg-specific T cells, which are induced/expanded in susceptible subjects and spontaneous disease models, likely because of defective immune regulation^1,50,51^, play an important role in islet destruction and T1D onset. Immune regulatory defects and T1D disease susceptibility in humans and mouse models have been linked to multiple genetic loci including that encoding CTLA-4^52^. Splice variants of CTLA-4 have been identified as potential risk factors contributing to the development of T1D^2^. While CTLA4 is considered as the master checkpoint receptor on T cells, other receptors such as PD1 and BTLA also play important roles in maintaining peripheral tolerance^6,8,53,54^. Checkpoint inhibitor therapy for cancers show that interfering with their functions, CTLA4 and PD1 functions particularly, can effectively break the T cell tolerance and treat many cancers^55^. Recent reports show that, although these therapies are effective against cancers, they produce autoimmune manifestations that include T1D and rheumatic diseases^31-33,56,57^, substantiating their role in peripheral self-tolerance. Therefore, these checkpoint receptors can, individually or in combinations, be targeted for promoting BcAg-specific T cell tolerance and preventing/suppressing spontaneous autoimmunity in T1D. Importantly, inhibition of T cell response through checkpoint receptors and inducing antigen specific tolerance requires delivering active negative signals through them in conjunction with TCR engagement^53,58-61^. Therefore, induction of BcAg-specific tolerance is possible because targeting CTLA4, PD1 and/or BTLA on T cells from professional antigen presenting cells such as DCs and enhancing the negative signaling strength occurs concurrently with the engagement of specific TCRs on T cells.

DCs are considered the most effective APCs and the only type of cells that can activate naïve T cells^62,63^. Hence, engineering the DCs to ensure maintenance of their tolerogenic function, even under pro-inflammatory conditions, is critical for establishing antigen specific T cell tolerance to prevent and treat T1D. In this regard, DCs that are selectively deleted of costimulatory molecules or ectopically expressing cytokine and non-cytokine factors have shown the ability to induce T cell tolerance^64-66^. We also reported that co-stimulation by CD80, preferential ligand of CTLA4 in terms of binding avidity, results in the generation of IL-10 dependent TGF-β1+ regulatory T cells ^35-39^. We reported that enhancing CTLA4 agonist strength on mature DC surface could lead to the dominant engagement of this repressor-receptor on T cells upon antigen presentation and induce antigen specific tolerance and prevent/suppress autoimmune thyroiditis and T1D^38,39^. Recently, using antigen immunization models, we demonstrated that lentiviral transduction approach can be employed to generate tDCs that are exogenously expressing selective ligands for multiple T cell repressor receptors (viz: CTLA4, PD1 and BTLA) and these DCs, particularly B7.1wa-DCs and multiligand-DCs as described in this study, can hyper-activate T cell inhibitory pathways individually or in combination during antigen presentation and induce antigen specific immune tolerance^40^. Our current study shows that treatments using B7.1wa-DCs and multiligand-DCs effectively induce BcAg tolerance and protects NOD mice from T1D.

Our current study shows that tDC, B7.1wa-DC and multiligand-DC, treatments promote protection from β-cell destruction and prevent T1D for a significant period when treated at pre-diabetic stage. These treatments also reverse the elevated blood glucose levels and improves the islet function significantly, when the treatment was initiated at an early-hyperglycemic stage. These observations suggest that a well-designed tDC approach such as the one described here has the potential to prevent and treat T1D in human subjects. Importantly, our observation that multiligand-DCs are more effective than B7.1wa-DCs in promoting protection from T1D in NOD mice suggests that evidence-based approaches for targeting multiple pathways is important for achieving effective and long-lasting T cell tolerance. Of note, our previous report also showed that while B7.1wa-DCs are more efficient in inducing Treg response and immune tolerance than the other checkpoint-ligand expressing DCs such PD-L1-DCs and HVEM-CRD1-DCs, and treatment using multiligand-DCs produced the most effective overall immune modulation^40^. Further, our previous reports have consistently demonstrated that enhanced engagement of CTLA4 during antigen presentation by DCs or autoimmune target tissue can produce immune tolerance and protection from autoimmunity, in both spontaneous and immunization-induced models^38,39^. However, the current study and our recent report^40^ demonstrates that concurrent engagement of other checkpoint receptors along with CTLA4 could enhance the tolerogenic ability of CTLA4 targeting DCs to achieve longer-lasting protection from spontaneous autoimmune manifestations.

Antigen specificity of the T cell tolerance induced by BcAg-pulsed B7.1wa- and multiligand-DCs is evident from different observations of this study. Not only the recipients of tDC-activated BDC2.5 T cells, but also those that received tDC-activated WT T cells failed to develop hyperglycemia. Moreover, challenge of immune cells from tDC treated mice using BcAg, but not anti-CD3 antibody, showed significantly diminished proliferation. Further, the release of large amounts of immune regulatory cytokines occurred primarily when the cells were challenged with BcAg. Importantly, our earlier study using animals that were immunized with ovalbumin or mouse-thyroglobulin^40^ has shown a profound increase in Foxp3+ T cells when treated with B7.1wa-DCs and multiligand-DCs. The current study using a spontaneous autoimmune diabetic model also showed significantly higher frequencies of Foxp3+ cells in ligand expressing DC treated mice compared to control-DC recipients.

Overall, our observations show that powerful tAPCs for T1D therapy can be generated by engineering the DCs to express the selective ligands of multiple T cell checkpoint receptors simultaneously, and this could be an attractive and effective approach for inducing BcAg specific T cell tolerance. Moreover, our previous report^40^ and the current study demonstrating the ability of these engineered DCs, multi-ligand DCs particularly, to affect multiple aspects of immune function, viz: higher Treg frequencies and immune regulatory cytokine responses, and lower pro-inflammatory IFNγ and IL17 responses against self-antigens suggest that such tDCs could have therapeutic value not only in T1D, but also can be exploited to treat other autoimmune diseases. Importantly, efficient engineering of the DCs for constitutive overexpression of multiple T cell inhibitory ligands, as described in this study, makes them less susceptible to changes, compared to previously employed immature DCs, in the functional properties upon in vivo therapeutic delivery; hence is desired for clinical translation for treating T1D.

## Author contribution

R.G. researched and analyzed data and edited the manuscript, S.K. researched data, N.P. researched data, G.L. researched data and C.V. designed experiments, researched and analyzed data, and wrote/edited manuscript.

## Data availability

The datasets generated during and/or analyzed during the current study are available from the corresponding author on reasonable request.

## Resource availability

The cDNA vectors generated during the current study is available from the corresponding author on reasonable request.

## Footnote

This work was supported by unrestricted research funds from MUSC and National Institutes of Health (NIH) grants R21AI069848, R01AI073858 and R21AI133798 to C.V. C.V. is the guarantor of this work and, as such, has full access to all the data in the study and takes responsibility for the integrity of data and accuracy of the data analysis. The authors are thankful to Cell and Molecular Imaging, Pathology, immune monitoring and discovery, and flow cytometry cores of MUSC for the histology service, microscopy, FACS and multiplex assay instrumentation support.

## References

1 Kukreja, A. et al. Multiple immuno-regulatory defects in type-1 diabetes. J Clin Invest109, 131–140, doi:10.1172/JCI13605 (2002).

2 Ueda, H. et al. Association of the T-cell regulatory gene CTLA4 with susceptibility to autoimmune disease. Nature 423, 506–511, doi:10.1038/nature01621 (2003).

3 Wicker, L. S. et al. Type 1 diabetes genes and pathways shared by humans and NOD mice. J Autoimmun 25 Suppl, 29–33, doi:10.1016/j.jaut.2005.09.009 (2005).

4 Zhang, Q. & Vignali, D. A. Co-stimulatory and Co-inhibitory Pathways in Autoimmunity. Immunity 44, 1034–1051, doi:10.1016/j.immuni.2016.04.017 (2016).

5 McGrath, M. M. & Najafian, N. The role of coinhibitory signaling pathways in transplantation and tolerance. Front Immunol 3, 47, doi:10.3389/fimmu.2012.00047 (2012).

6 Schildberg, F. A., Klein, S. R., Freeman, G. J. & Sharpe, A. H. Coinhibitory Pathways in the B7-CD28 Ligand-Receptor Family. Immunity 44, 955–972,doi:10.1016/j.immuni.2016.05.002 (2016).

7 Chen, L. & Flies, D. B. Molecular mechanisms of T cell co-stimulation and co-inhibition. Nat Rev Immunol 13, 227–242, doi:10.1038/nri3405 (2013).

8 Watanabe, N. et al. BTLA is a lymphocyte inhibitory receptor with similarities to CTLA-4 and PD-1. Nat Immunol 4, 670–679, doi:10.1038/ni944 (2003).

9 Wells, A. D., Walsh, M. C., Bluestone, J. A. & Turka, L. A. Signaling through CD28 and CTLA-4 controls two distinct forms of T cell anergy. The Journal of clinical investigation 108, 895–903, doi:10.1172/JCI13220 (2001).

10 Munn, D. H., Sharma, M. D. & Mellor, A. L. Ligation of B7-1/B7-2 by human CD4+ T cells triggers indoleamine 2,3-dioxygenase activity in dendritic cells. J Immunol 172, 4100–4110 (2004).

11 Greenwald, R. J. et al. CTLA-4 regulates cell cycle progression during a primary immune response. Eur J Immunol 32, 366–373, doi:10.1002/1521-4141(200202)32:2<366::AID-IMMU366>3.0.CO;2-5 (2002).

12 Vanasek, T. L., Khoruts, A., Zell, T. & Mueller, D. L. Antagonistic roles for CTLA-4 and the mammalian target of rapamycin in the regulation of clonal anergy: enhanced cell cycle progression promotes recall antigen responsiveness. J Immunol 167, 5636–5644 (2001).

13 Tang, Q. et al. Cutting edge: CD28 controls peripheral homeostasis of CD4+CD25+ regulatory T cells. J Immunol 171, 3348–3352 (2003).

14 Salomon, B. et al. B7/CD28 costimulation is essential for the homeostasis of the CD4+CD25+ immunoregulatory T cells that control autoimmune diabetes. Immunity 12, 431–440 (2000).

15 Tang, Q. et al. Distinct roles of CTLA-4 and TGF-beta in CD4+CD25+ regulatory T cell function. Eur J Immunol 34, 2996–3005, doi:10.1002/eji.200425143 (2004).

16 Takahashi, T. et al. Immunologic self-tolerance maintained by CD25(+)CD4(+) regulatory T cells constitutively expressing cytotoxic T lymphocyte-associated antigen 4. J Exp Med 192, 303–310 (2000).

17 Wing, K. et al. CTLA-4 control over Foxp3+ regulatory T cell function. Science 322, 271–275, doi:10.1126/science.1160062 (2008).

18 Matheu, M. P. et al. Imaging regulatory T cell dynamics and CTLA4-mediated suppression of T cell priming. Nat Commun 6, 6219, doi:10.1038/ncomms7219 (2015).

19 Qureshi, O. S. et al. Trans-endocytosis of CD80 and CD86: a molecular basis for the cell-extrinsic function of CTLA-4. Science 332, 600–603, doi:10.1126/science.1202947 (2011).

20 Khattri, R., Auger, J. A., Griffin, M. D., Sharpe, A. H. & Bluestone, J. A. Lymphoproliferative disorder in CTLA-4 knockout mice is characterized by CD28-regulated activation of Th2 responses. J Immunol 162, 5784–5791 (1999).

21 Tivol, E. A. et al. Loss of CTLA-4 leads to massive lymphoproliferation and fatal multiorgan tissue destruction, revealing a critical negative regulatory role of CTLA-4. Immunity 3, 541–547 (1995).

22 Chemnitz, J. M., Parry, R. V., Nichols, K. E., June, C. H. & Riley, J. L. SHP-1 and SHP-2 associate with immunoreceptor tyrosine-based switch motif of programmed death 1 upon primary human T cell stimulation, but only receptor ligation prevents T cell activation. J Immunol 173, 945–954 (2004).

23 Liang, S. C. et al. Regulation of PD-1, PD-L1, and PD-L2 expression during normal and autoimmune responses. Eur J Immunol 33, 2706–2716, doi:10.1002/eji.200324228 (2003).

24 Nishimura, H. et al. Autoimmune dilated cardiomyopathy in PD-1 receptor-deficient mice. Science 291, 319–322, doi:10.1126/science.291.5502.319 (2001).

25 Gonzalez, L. C. et al. A coreceptor interaction between the CD28 and TNF receptor family members B and T lymphocyte attenuator and herpesvirus entry mediator. Proc Natl Acad Sci U S A 102, 1116–1121, doi:10.1073/pnas.0409071102 (2005).

26 Croft, M. The evolving crosstalk between co-stimulatory and co-inhibitory receptors: HVEM-BTLA. Trends Immunol 26, 292–294, doi:10.1016/j.it.2005.03.010 (2005).

27 Cai, G. et al. CD160 inhibits activation of human CD4+ T cells through interaction with herpesvirus entry mediator. Nat Immunol 9, 176–185, doi:10.1038/ni1554 (2008).

28 Seidel, J. A., Otsuka, A. & Kabashima, K. Anti-PD-1 and Anti-CTLA-4 Therapies in Cancer: Mechanisms of Action, Efficacy, and Limitations. Front Oncol 8, 86, doi:10.3389/fonc.2018.00086 (2018).

29 Chan, T. A., Wolchok, J. D. & Snyder, A. Genetic Basis for Clinical Response to CTLA-4 Blockade in Melanoma. N Engl J Med 373, 1984, doi:10.1056/NEJMc1508163 (2015).

30 Boussiotis, V. A. Molecular and Biochemical Aspects of the PD-1 Checkpoint Pathway. N Engl J Med 375, 1767–1778, doi:10.1056/NEJMra1514296 (2016).

31 Ramos-Casals, M. et al. Immune-related adverse events of checkpoint inhibitors. Nat Rev Dis Primers 6, 38, doi:10.1038/s41572-020-0160-6 (2020).

32 Quandt, Z., Young, A. & Anderson, M. Immune checkpoint inhibitor diabetes mellitus: a novel form of autoimmune diabetes. Clinical and experimental immunology 200, 131–140, doi:10.1111/cei.13424 (2020).

33 Braaten, T. J. et al. Immune checkpoint inhibitor-induced inflammatory arthritis persists after immunotherapy cessation. Ann Rheum Dis 79, 332–338, doi:10.1136/annrheumdis-2019-216109 (2020).

34 Melissaropoulos, K. et al. Rheumatic Manifestations in Patients Treated with Immune Checkpoint Inhibitors. Int J Mol Sci 21, doi:10.3390/ijms21093389 (2020).

35 Vasu, C., Prabhakar, B. S. & Holterman, M. J. Targeted CTLA-4 engagement induces CD4+CD25+CTLA-4high T regulatory cells with target (allo)antigen specificity. Journal of immunology 173, 2866–2876 (2004).

36 Vasu, C., Gorla, S. R., Prabhakar, B. S. & Holterman, M. J. Targeted engagement of CTLA-4 prevents autoimmune thyroiditis. International immunology 15, 641–654 (2003).

37 Perez, N. et al. Preferential costimulation by CD80 results in IL-10-dependent TGF-beta1(+) -adaptive regulatory T cell generation. J Immunol 180, 6566–6576 (2008).

38 Li, R. et al. Enhanced engagement of CTLA-4 induces antigen-specific CD4+CD25+Foxp3+ and CD4+CD25-TGF-beta 1+ adaptive regulatory T cells. Journal of immunology 179, 5191–5203 (2007).

39 Karumuthil-Melethil, S. et al. Dendritic cell-directed CTLA-4 engagement during pancreatic beta cell antigen presentation delays type 1 diabetes. J Immunol 184, 6695–6708, doi:10.4049/jimmunol.0903130 (2010).

40 Gudi, R. R., Karumuthil-Melethil, S., Perez, N., Li, G. & Vasu, C. Engineered Dendritic Cell-Directed Concurrent Activation of Multiple T cell Inhibitory Pathways Induces Robust Immune Tolerance. Sci Rep 9, 12065, doi:10.1038/s41598-019-48464-y (2019).

41 Korn, T. et al. Myelin-specific regulatory T cells accumulate in the CNS but fail to control autoimmune inflammation. Nat Med 13, 423–431, doi:nm1564 [pii] 10.1038/nm1564 (2007).

42 Sofi, M. H. et al. Polysaccharide A-Dependent Opposing Effects of Mucosal and Systemic Exposures to Human Gut Commensal Bacteroides fragilis in Type 1 Diabetes. Diabetes 68, 1975–1989, doi:10.2337/db19-0211 (2019).

43 Karumuthil-Melethil, S. et al. TLR2-and Dectin 1-associated innate immune response modulates T-cell response to pancreatic beta-cell antigen and prevents type 1 diabetes. Diabetes 64, 1341–1357, doi:10.2337/db14-1145 (2015).

44 Karumuthil-Melethil, S., Gudi, R., Johnson, B. M., Perez, N. & Vasu, C. Fungal beta-glucan, a Dectin-1 ligand, promotes protection from type 1 diabetes by inducing regulatory innate immune response. J Immunol 193, 3308–3321, doi:10.4049/jimmunol.1400186 (2014).

45 Throm, R. E. et al. Efficient construction of producer cell lines for a SIN lentiviral vector for SCID-X1 gene therapy by concatemeric array transfection. Blood 113, 5104–5110, doi:10.1182/blood-2008-11-191049 (2009).

46 Gudi, R. et al. Complex dietary polysaccharide modulates gut immune function and microbiota, and promotes protection from autoimmune diabetes. Immunology 157, 70–85, doi:10.1111/imm.13048 (2019).

47 Sofi, M. H. et al. pH of drinking water influences the composition of gut microbiome and type 1 diabetes incidence. Diabetes 63, 632–644, doi:db13-0981 [pii] 10.2337/db13-0981 (2014).

48 Zheng, P., Wu, Y., Guo, Y., Lee, C. & Liu, Y. B7-CTLA4 interaction enhances both production of antitumor cytotoxic T lymphocytes and resistance to tumor challenge. Proceedings of the National Academy of Sciences of the United States of America 95, 6284–6289, doi:10.1073/pnas.95.11.6284 (1998).

49 Sedy, J. R. et al. B and T lymphocyte attenuator regulates T cell activation through interaction with herpesvirus entry mediator. Nat Immunol 6, 90–98, doi:10.1038/ni1144 (2005).

50 Pugliese, A. Autoreactive T cells in type 1 diabetes. J Clin Invest 127, 2881–2891, doi:10.1172/JCI94549 (2017).

51 Zoka, A. et al. Altered immune regulation in type 1 diabetes. Clin Dev Immunol 2013, 254874, doi:10.1155/2013/254874 (2013).

52 Pociot, F. & McDermott, M. F. Genetics of type 1 diabetes mellitus. Genes Immun 3, 235–249, doi:10.1038/sj.gene.6363875 (2002).

53 Walunas, T. L., Bakker, C. Y. & Bluestone, J. A. CTLA-4 ligation blocks CD28-dependent T cell activation. J Exp Med 183, 2541–2550 (1996).

54 Freeman, G. J. et al. Engagement of the PD-1 immunoinhibitory receptor by a novel B7 family member leads to negative regulation of lymphocyte activation. J Exp Med 192, 1027–1034 (2000).

55 Robert, C. A decade of immune-checkpoint inhibitors in cancer therapy. Nat Commun 11, 3801, doi:10.1038/s41467-020-17670-y (2020).

56 Lidar, M. et al. Rheumatic manifestations among cancer patients treated with immune checkpoint inhibitors. Autoimmun Rev 17, 284–289, doi:10.1016/j.autrev.2018.01.003 (2018).

57 Suarez-Almazor, M. E., Kim, S. T., Abdel-Wahab, N. & Diab, A. Review: Immune-Related Adverse Events With Use of Checkpoint Inhibitors for Immunotherapy of Cancer. Arthritis Rheumatol 69, 687–699, doi:10.1002/art.40043 (2017).

58 Hwang, K. W. et al. Cutting edge: targeted ligation of CTLA-4 in vivo by membrane-bound anti-CTLA-4 antibody prevents rejection of allogeneic cells. Journal of immunology 169, 633–637, doi:10.4049/jimmunol.169.2.633 (2002).

59 Griffin, M. D. et al. Blockade of T cell activation using a surface-linked single-chain antibody to CTLA-4 (CD152). Journal of immunology 164, 4433–4442, doi:10.4049/jimmunol.164.9.4433 (2000).

60 Chai, J. G., Vendetti, S., Amofah, E., Dyson, J. & Lechler, R. CD152 ligation by CD80 on T cells is required for the induction of unresponsiveness by costimulation-deficient antigen presentation. J Immunol 165, 3037–3042 (2000).

61 Fife, B. T., Griffin, M. D., Abbas, A. K., Locksley, R. M. & Bluestone, J. A. Inhibition of T cell activation and autoimmune diabetes using a B cell surface-linked CTLA-4 agonist. J Clin Invest 116, 2252–2261, doi:10.1172/JCI27856 (2006).

62 Turley, S. J. Dendritic cells: inciting and inhibiting autoimmunity. Curr Opin Immunol 14, 765–770 (2002).

63 Bayry, J. et al. Dendritic cells and autoimmunity. Autoimmun Rev 3, 183–187, doi:10.1016/S1568-9972(03)00104-6 (2004).

64 Liang, X. et al. Administration of dendritic cells transduced with antisense oligodeoxyribonucleotides targeting CD80 or CD86 prolongs allograft survival. Transplantation 76, 721–729, doi:10.1097/01.TP.0000076470.35404.49 (2003).

65 Prechtel, A. T., Turza, N. M., Theodoridis, A. A. & Steinkasserer, A. CD83 knockdown in monocyte-derived dendritic cells by small interfering RNA leads to a diminished T cell stimulation. J Immunol 178, 5454–5464 (2007).

66 Besche, V. et al. Dendritic cells lentivirally engineered to overexpress interleukin-10 inhibit contact hypersensitivity responses, despite their partial activation induced by transduction-associated physical stress. J Gene Med 12, 231–243, doi:10.1002/jgm.1436 (2010).

