## supplemental figures for "Activation of T cell checkpoint pathways during β-cell antigen presentation by engineered dendritic cells promotes protection of non-obese diabetic mice from type 1 diabetes"

Supplemental fig. 1

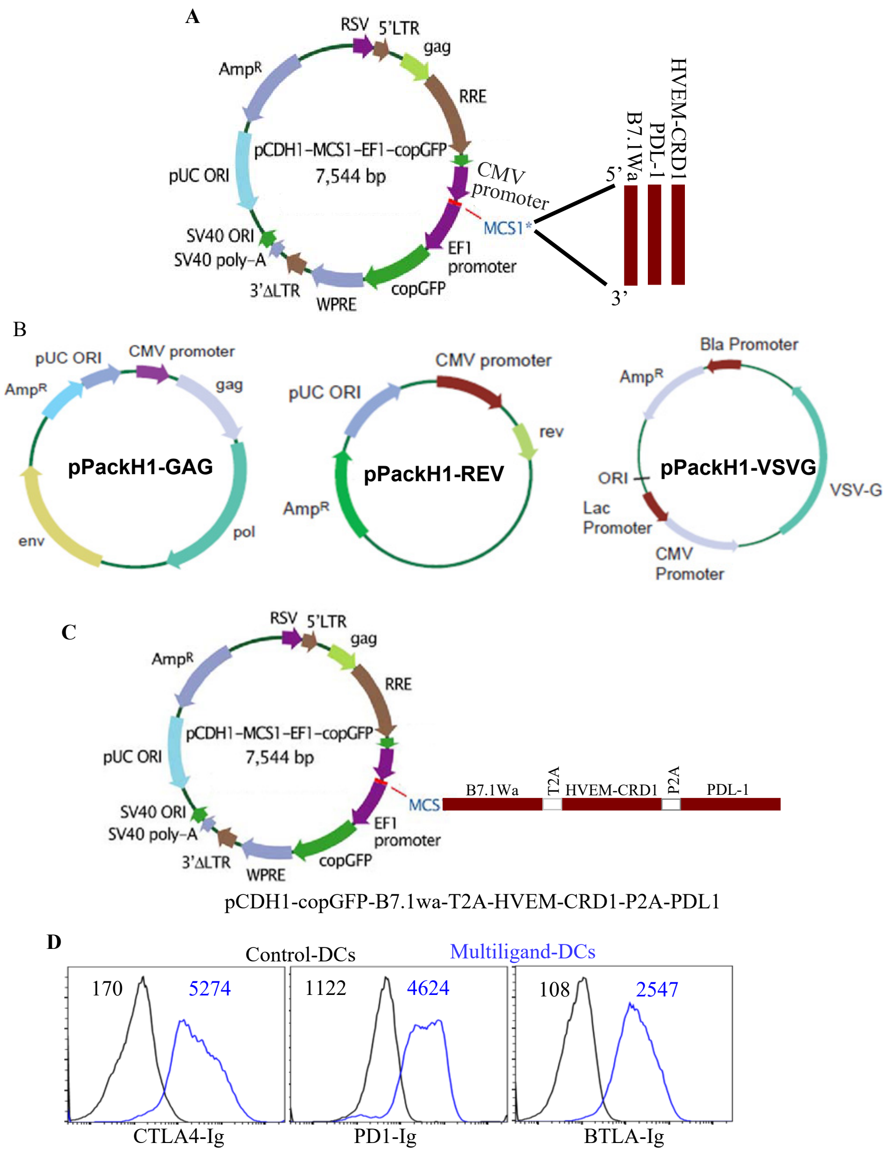


**Supplemental Fig. 1: Lentiviral system and the constructs used in this study. A)** T cell negative regulatory ligands were cloned individually without fluorescent tags under the CMV promoter of a 3$^{rd}$ generation lentiviral vector from SBI Inc. In these vectors, GFP is expressed under a different (EF1) promoter. Unmodified vector was used as control. **B)** Shows packaging vectors originally purchased as a pool from SBI were separated, propagated and used for generating lentivirus in HEK293T cells or GFPG cells. **C)** In some experiments, a multiligand-cDNA construct was used for expressing multiple ligands from the same vector. Individual cDNAs were separated by self-cleaving peptides as indicated. **D)** Functional ligand expression levels on control- and multiligand-DCs (generated using the vector constructs shown in panel C) were determined after incubating with soluble receptors (CTLA4-Ig, PD1-Ig or BTLA-Ig), followed by PE-labeled Fab(2) fragment of anti-IgG (Fc specific) Ab and testing by FACS. Histogram overlay graphs (representative of 3 independent experiments) showing ligand specific staining of control virus transduced and multiligand virus transduced DCs along with MFI values are shown.
